# Topological evolution of sprouting vascular networks: from day-by-day analysis to general growth rules

**DOI:** 10.1101/2023.09.02.555959

**Authors:** Katarzyna O. Rojek, Antoni Wrzos, Stanisław Żukowski, Michał Bogdan, Maciej Lisicki, Piotr Szymczak, Jan Guzowski

**Affiliations:** Institute of Physical Chemistry, Polish Academy of Sciences, Warsaw, Poland; Institute of Theoretical Physics, Faculty of Physics, University of Warsaw, Warsaw, Poland; Laboratoire Matière et Systèmes Complexes (MSC), UMR 7057, CNRS & Université Paris Cité, Paris, France

**Keywords:** angiogenesis, bead sprouting assay, image analysis

## Abstract

Engineering tissues with an embedded vasculature of well-controlled topology remains one of the basic problems in biofabrication. Still, little is known about the evolution of topological characteristics of vascular networks over time. Here, we perform a high-throughput day-by-day analysis of tens of microvasculatures that sprout from endothelial-cell coated micrometric beads embedded in an external fibrin gel. We use the bead-assays to systematically analyze (i) ‘macroscopic’ observables such as the overall length and area of the sprouts, (ii) ‘microscopic’ observables such as the lengths of segments or the branching angles and their distributions, as well as (iii) general measures of network complexity such as the average number of bifurcations per branch. We develop a custom angiogenic image analysis toolkit and track the evolution of the networks for at least 14 days of culture under various conditions, e.g., in the presence of fibroblasts or with added endothelial growth factor (VEGF). We find that the evolution always consists of three stages: (i) an inactive stage in which cells remain bound to the beads, (ii) a sprouting stage in which the sprouts rapidly elongate and bifurcate, and (iii) the maturation stage in which the growth slows down. We show that higher concentrations of VEGF lead to an earlier onset of sprouting and to a higher number of primary branches, yet without significantly affecting the speed of growth of the individual sprouts. We find that the mean branching angle is weakly dependent on VEGF and typically in the range of 60-75 degrees suggesting that, by comparison with the available Laplacian growth models, the sprouts tend to follow local VEGF gradients. Finally, we observe an exponential distribution of segment lengths, which we interpret as a signature of stochastic branching at a constant bifurcation rate (per unit branch length). Our results, due to high statistical relevance, may serve as a benchmark for predictive models and reveal how the external means of control, such as VEGF concentration, could be used to control the morphology of the vascular networks. We provide guidelines for the fabrication of optimized microvasculatures with potential applications in drug testing or regenerative medicine.

## INTRODUCTION

Microvascular tissue engineering is an emerging subfield in tissue engineering that focuses on the development of living capillary networks ^1-3^ and vascularized microtissues for applications in drug testing ^4-9^, regenerative medicine ^10-13^, and general biofabrication ^3^. It is known that most types of engineered tissue constructs, exceeding in size the so-called diffusion limit (appr. 1 mm), eventually suffer from hypoxia if deprived of microvasculature ^14^. Accordingly, engineering of viable tissues at all scales necessarily requires the incorporation of an embedded microvasculature. However, thus far, despite a large body of literature regarding microvascular networks observed both *in vivo* ^15-17^ and *in vitro* ^6, 16^, the general rules governing the development of branching capillary networks in terms of their dynamic topology, e.g., bifurcation frequency, bifurcation angles, lacunarity, etc., remain obscure. Here, we address this problem by tracking the evolution of dozens of microvascular networks day-by-day, all cultured in similar, controlled conditions. We develop custom image analysis tools to extract morphological/topological characteristics of the networks in terms of statistical distributions of several observables (segment lengths, branching angles, etc.) which allows us to propose basic rules governing the development of sprouting vascular networks.

In this work, we use the so-called bead-sprouting assays in which the capillary networks ‘sprout’ from the endothelial cell-coated microbeads suspended in an external 3D hydrogel matrix. We exploit the sprouting networks as a convenient model system for studying microvascular dynamics with potential future applications, e.g., in drug testing. However, in further perspective, as we also detail below, such modular microvasculatures could prove useful as building blocks of larger interconnected vascular networks for applications in regenerative medicine or bioprinting.

Endothelial cells (ECs), the main constituents of vessel walls, are known to spontaneously form vascular network-like structures when embedded in the external extracellular matrix (ECM). Vasculogenesis occurs naturally, e.g., during embryonic development but has also been exploited as a route towards self-assembly of microvascular structures *in vitro* including perfusable vasculature-on-chip systems ^4, 8^. Typically, in such approaches, ECs are initially dispersed randomly in the external ECM-like hydrogel and, with time, proliferate and migrate to form a network of microvessels.

Alternatively, the formation of capillary networks *in vitro* can also be achieved via angiogenesis, that is the outgrowth or ‘sprouting’ of new vessels from a pre-existing vasculature or an EC-monolayer. In such a case, tree-like capillary networks emerge via sequential branching of the sprouting capillaries supported by cell migration along the sprouts ^6, 18-20^.

From the point of view of tissue engineering, it would be optimal to exploit both vasculogenesis and angiogenesis at the same time, since both strategies have their unique advantages. In vasculogenesis, proliferation and free migration of cells rapidly lead to the development of an interconnected bulk vasculature. In angiogenesis, growth occurs mainly at the tips of the sprouts, resulting in slower percolation of the bulk ECM. However, the angiogenic sprouting naturally leads to a spatially more organized, hierarchical vasculature, potentially useful in terms of targeted transport of nutrients, efficient perfusion and easier in integration with the external channels enabling organ-on-chip applications ^21^. Accordingly, to achieve organized networks at possibly short culture times, one would like to combine the advantages of both vasculogenesis and angiogenesis. A convenient approach towards this goal is the use of endothelial cell carriers that can be dispersed throughout the ECM to serve as sprouting ‘seeds’. In this strategy, the multiple sprouting microvasculatures are supposed to eventually interconnect to provide a fully percolated mesoscale network. The method has been originally proposed by Stegemann, Putnam and co-workers ^22^ and more recently picked up by others ^23, 24^. EC-encapsulating fibrin beads ^22^ or EC-coated synthetic hydrogel beads ^24^ were used as the sprouting seeds. At late culture times, typically around or after day 7, neighboring sprouting microvasculatures were observed to form interconnections. However, the dynamic processes of network formation were not studied in detail.

In fact, even in the case of an isolated bead-sprouting network, the growth processes remain poorly understood. Quite surprisingly, despite the rich literature on the biological complexity of angiogenic sprouts, e.g., differentiation of the stalk and tip cells ^24-27^ and theoretical models of capillary network formation ^28-31^, little is known about the different stages of development of the tree-like endothelial networks. In particular, the dynamical changes in network connectivity, fractal dimension, lacunarity and overall network complexity and topology have not been studied in detail. Also, the impact of external biochemical and/or biomechanical cues on the angiogenic dynamics remains obscure. In fact, the available literature has mostly focused on the analysis of global parameters of the sprouting networks, such as the final total length of the sprouts, total area, etc. ^32-34^, and typically, at late times or at maximum 3 time points, which can be used to estimate the overall trends in growth ^32, 33^ yet say little about the evolving topology. From a tissue-engineering point of view, the lack of knowledge about general rules governing the topological evolution of sprouting vascular networks limits further progress in biofabrication of viable tissues ^11, 35-37^.

Here, we make a first step towards establishing such general rules via performing high-throughput parallel bead-sprouting experiments and their careful and systematic analysis. We independently track the evolution of tens of sprouting vascular networks day-by-day over the course of at least 14 days. To this end, we additionally develop custom image analysis scripts which involve skeletonization of the network, branching-point detection and bifurcation angle measurement. We use the software to quantitatively analyze the evolution of (i) *global observables*, such as the overall length and area of the sprouts, (ii) *microscopic observables*, such as branch lengths and branching angles, and their distributions, as well as (iii) *general measures of network complexity* such as the average number of bifurcations per branch. We reveal that the evolution of the sprouting networks proceeds through three stages: (i) the ‘idle’ stage during which the cells proliferate only within the monolayer covering the bead, (ii) the exponential sprout growth stage, consisting of rapid sprout elongation and branching, and (iii) the maturation stage during which the growth slows down. We also study the impact of the presence of fibroblasts in the ECM (either as a monolayer on top of ECM or intermixed within ECM) and the concentration of the vascular endothelial growth factor (VEGF) in the culture media on the vascular growth dynamics.

We discuss our results in terms of the existing models of network formation and find that the results support a picture of a stochastically branching network. One of our primary findings is that at larger VEGF concentrations (25-50 ng/ml) the distribution of segment lengths (i.e. distances between the bifurcation points) becomes Poissonian. This observation remains in line with the available branching morphogenesis models ^38^. Interestingly, the characteristic length does not seem to depend on the VEGF concentration. We also find that the average values of the branching angle vary in the range of 60-75 degrees, i.e., close to (2π/5) x 180 = 72 degrees characteristic of the Laplacian growth models ^39-42^, which overall suggests that the tips follow the local gradients of the growth factor concentration. Collectively, our results, due to their high statistical relevance, may serve as a benchmark for predictive models and reveal how the external means of control, such as VEGF concentration, could be used to control the topology and morphology of the early microvascular networks.

## RESULTS

### Long-term time-lapse imaging of endothelial sprouting of single EC-coated beads

To study the morphogenesis of vascular networks, we examined the dynamics of EC sprouting form isolated EC-coated beads during the *in vitro* angiogenic process. We coated polystyrene microbeads with GFP-tagged human umbilical vein endothelial cells (HUVECs) (Figure 1A) and embedded them in 2.5 mg/ml fibrin hydrogel. The beads were seeded in 24 well plates, with one bead per well (Figure 1B) to exclude the potential impact of the presence of nearby beads on the directionality and/or growth dynamics of the angiogenic sprouts ^23^. Also, to provide possibly isotropic initial conditions, the beads were positioned centrally in the wells, at large well-to-bead size ratio *d*_well_/*d*_bead_ ≈ 60. During the assay, we observed several stages of the angiogenic process starting with the formation of the endothelial layer at the surface of the bead, followed by sprouting of the ECs from the layer, sprout elongation, bifurcation, and finally the formation of a characteristic dendritic star-like architecture. To characterize the evolution of the global properties of the network over time, we visualized the morphology of the angiogenic sprouts around each of the EC-coated beads using confocal microscopy. We acquired the images at 24 h time intervals over the period of 14 days to cover all stages of the angiogenic morphogenesis (Figure 1C). We placed the samples under a confocal microscope once per day for a short time to acquire an image of the EC-coated bead whereas in between the imaging sessions the sample was placed in the CO_2_ incubator. Such intermittent imaging sessions limited the time of exposure of the cells to the non-physiological conditions outside the CO_2_ incubator and as supported cell viability and extended their lifetime. We observed cells to remain viable for at least 14 days, and even up to 20 days in some cases (Supp. Figure 1).

**Figure 1.**
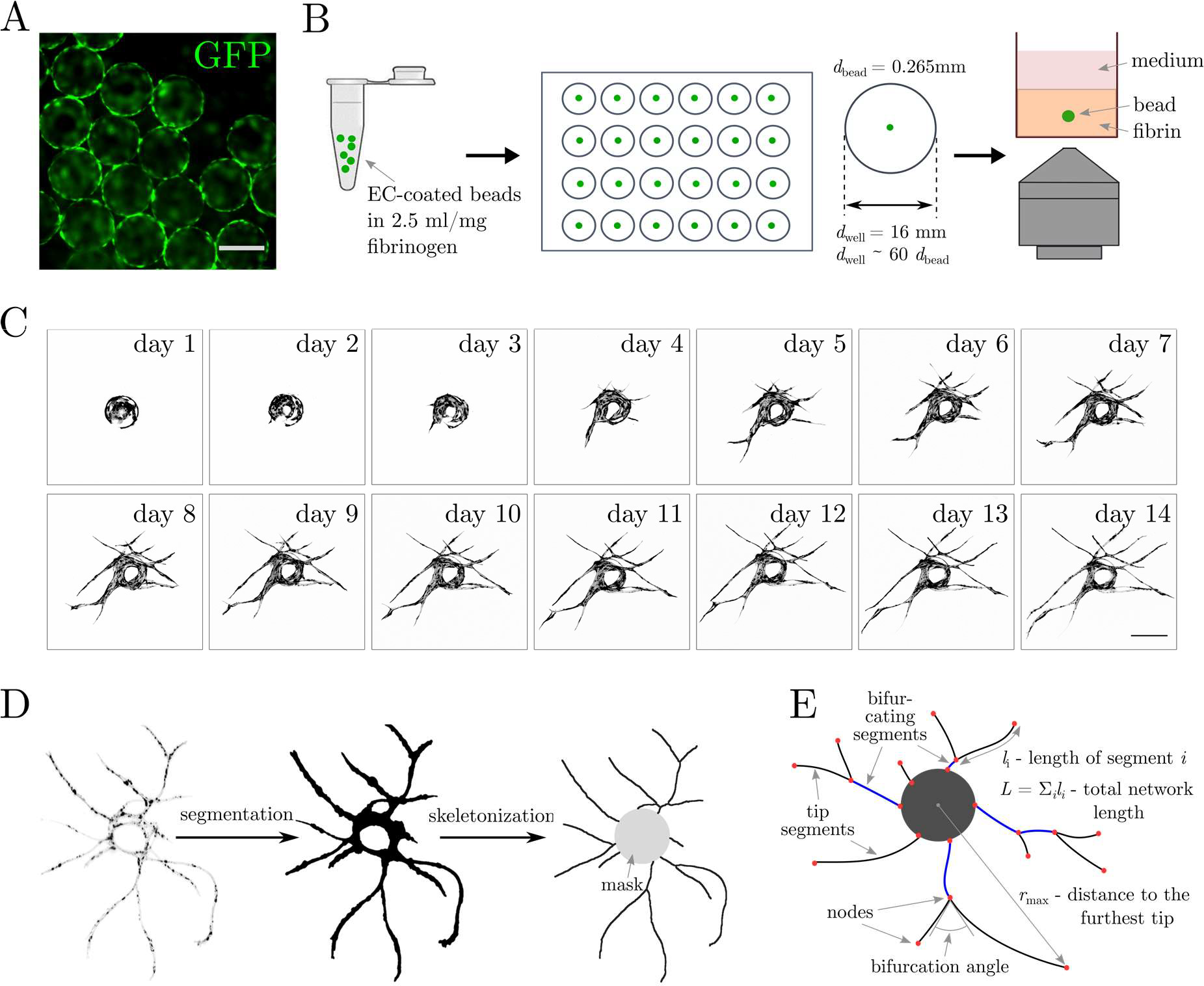
Long-term time-lapse imaging of endothelial sprouting of single EC-coated beads: experimental design and image analysis workflow. **(A)** Polystyrene beads coated with GFP-transduced HUVECs. **(B)** Scheme illustrating the experimental workflow. Cell-coated beads are resuspended in fibrin solution, seeded one bead per well in a 24-well plate and imaged using confocal microscopy. **(C)** Representative confocal microscopy images of a single HUVEC-coated bead acquired at one day intervals for 14 consecutive days. **(D)** Image processing workflow. From left to right: segmentation, skeletonization and conversion to a graph. **(E)** Schematic representation of the basic metrics used to characterize the networks. The scale bars in (A, C) are 250 μm.

The proposed approach allowed us to track the evolution of a high number of sprouting networks in parallel. From a practical point of view, the setup did not require any additional medium supply; the medium could be easily changed under a laminar hood between imaging sessions. Our day-by-day analysis goes beyond the previous studies which typically focused either on a single culture time point, e.g., day 7 or day 14 ^32-34^, or studied the EC sprout outgrowth during a relatively short window of time, usually limited to 1-2 days, at high image acquisition frequency ^16, 25^. The latter approach often suffered from photobleaching and other light-exposure related issues ^43^.

### Image analysis

To efficiently analyze the large amount of generated data, we have developed an automated image processing tool. The software overlays confocal microscopy images of a given sample acquired at multiple time points to allow tracking the evolution of sprouts around each given bead. The program extracts the morphology of a sprouting network directly from the experimental image and produces a database of the measured metrics (bifurcation and termination points, segment lengths, branching angles, etc.) which are further analyzed statistically (Figure 1D, E). We use Python as the programming language which offers open-source image processing libraries and significantly simplifies the code ^44-46^.

Our image processing workflow starts with image segmentation. The software detects the EC-coated polystyrene bead, defines a corresponding circular ‘mask’ and identifies the sprouts. Next, it calculates the projected area of the cells inside the mask (*A*_c_, the area of the cells covering the bead) and outside the mask (*A*, the total area of the sprouts) as well as the total length of the vascular network (*L*) and the mean width of the sprout (*λ* = *A*/*L*). Then it performs skeletonization of the previously segmented image and saves it as a custom graph class (Figure 1D). The nodes of the graph are identified as one of the following: (i) a branching point, (ii) a base of a sprout (the point of contact with the mask), or (iii) a sprout tip. The sprout *segments* are accordingly identified as the parts of sprouts contained between two neighboring nodes. In particular, the software separately classifies the *tip segments*, that is the segments terminating with a sprout tip, i.e., non-bifurcating ones, and the *bifurcating segments*. In the following, we use subscripts *tip* or *bif* to distinguish between the two types of segments.

Next, the program calculates the radial coordinates of the tips and finds their maximum (*r*_max_). It also finds the number of primary branches (*N*_pb_), i.e., the segments in direct contact with the central bead, and the number of tips (*N*_tip_). The complexity of the vascular network is assessed by computing the average number of generations *G* associated with each of the tree-like subnetworks originating from a primary branch, that is, *G* = 1+log_2_(*N*_tip_/*N*_pb_).

Finally, our code measures the bifurcation angles (*ϕ*), which, to our best knowledge, have not yet been quantified previously in the context of microvascular networks (Figure 1E). To this end, in particular, the software applies a filter which allows one to distinguish (in most cases) between bifurcations and anastomosis events and focuses only on the former.

### The interstitial distribution of fibroblasts promotes endothelial sprouting of EC-coated beads

It is known that fibroblasts act as supporting cells and surround capillary-like structures, promoting the formation of stable vascular networks ^47^. Fibrinolytic activity of fibroblasts leads to remodeling and gradual degradation of the fibrin matrix ^47^. This enhances the diffusive transport of fibroblast-derived proangiogenic factors, such as vascular endothelial growth factor (VEGF), angiopoietin-1 and platelet-derived growth factor (PDGF), and, in general, has a supportive effect on the diffusion of the cell culture media throughout the fibrin matrix ^33^. In the first series of experiments, we aimed at examining the impact of the distribution of normal human dermal fibroblasts (NHDFs) within the fibrin matrix on the morphogenesis of the vascular network around an isolated EC-coated bead. NHDFs were either distributed in the bulk of the fibrin gel (the ‘intermixed’ case) or seeded at the gel-media interface forming a cellular monolayer (the ‘monolayer’ case), see Figure 2A and Supp. Figure 2. We observed that, in qualitative agreement with previous studies ^32, 33, 47^, distributing fibroblasts interstitially, promoted sprouting and led to faster network development as compared to the case of a monolayer. This was reflected in a significant increase in the total area *A* and length *L* of the vascular networks at all times, see Figures 2Ba and 2Bb. The difference was already apparent at day 5 of culture and gradually increased, reaching the maximum at day 14. Also, starting from day 7, the rate of growth of both *A* and *L* appeared approximately three times higher in the ‘intermixed’ case, with the measured values doubling those observed in the ‘monolayer’ case at day 14. The observed dynamics strongly correlated with the dynamics of the number of primary branches *N*_pb_, the number of tips *N*_tip_ and the distance to the furthest tip *r*_max_ (Figure 2Be-2Bg), despite somewhat smaller differences between the ‘intermixed’ and ‘monolayer’ configurations in these cases. Overall, in the ‘monolayer’ case the dynamics of *A, L*, and *N*_pb_, *N*_tip_, *r*_max_ significantly slowed down starting from day 6 or 7 (depending on the observable), whereas in the ‘intermixed’ case the pace of growth remained high until much later times, that is day 14 (for *A* and *L*) or day 11 (for *N*_pb_ and *N*_tip_). Considering the maximal distances *r*_max_ span by the networks, the ‘intermixed’ networks grew wider by roughly 200 μm as compared to the ‘monolayer’ networks. Interestingly, the difference emerged in a stepwise manner, with the jump occurring around days 7 and 8, and remained roughly unchanged at later times. Hence, it seems that distributing fibroblasts inside the ECM tends to extend the period of intensive sprout elongation. The complexity of the network, as measured by the number of branch generations *G*, remained slightly elevated in the ‘intermixed’ case, starting from day 8 (Figure 2Bh). This suggests that distributing NHDFs through the fibrin matrix leads to the formation of bigger (in terms of *L, A*) and slightly more branched (*G*) vascular networks. Finally, the number of cells covering the bead, as measured by *A*_c_ appeared not to be significantly affected by the fibroblast distribution (Figure 2Bd). The values measured in the ‘monolayer’ case remained around 10% higher than in the ‘intermixed’ case. The shortage of bead-coating cells in the ‘intermixed’ case could be related to the higher number of cells migrating from the bead towards the sprouts in this case. The average width of the sprouts (*λ*) was very similar in both cases starting from day 6, i.e., once the sprouting set off in both cases (Figure 2Bc).

**Figure 2.**
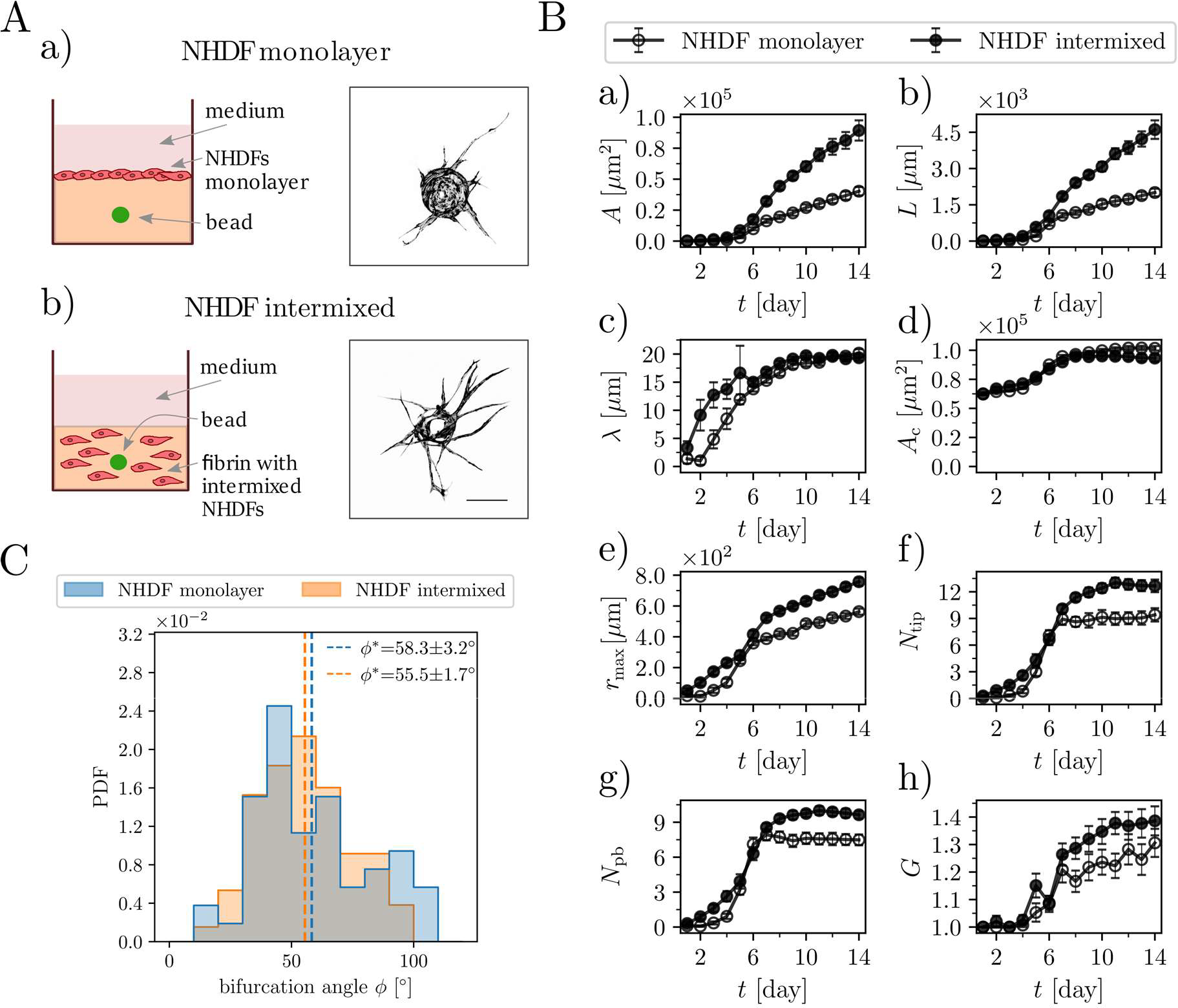
Distributing NHDFs through the matrix promotes endothelial sprouting of EC-coated beads. (A) Different seeding condition of NHDFs during the angiogenesis bead-sprouting assay – schematic drawings and representative images of single GFP-tagged HUVEC-coated beads. NHDFs were either seeded as (a) a monolayer on the top of a fibrin clot or (b) distributed through the fibrin matrix. Day 10 of culture. Scale bar 250 μm. **(B)** Morphometric analysis of (a) total area *A*, (b) total length *L*, (c) average sprout width *λ*, (d) area of the cell-coated bead *A*_*c*_, (e) distance to the furthest tip *r*_*max*,_ (f) number of tips *N*_*tip*_, (g) number of primary branches *N*_*pb*_ and (h) the average number of generations *G* per branch (*G =* log_2_(*N*_*tip*_*/N*_*pb*_)) of the sprouting capillary networks for 14 consecutive days. The numbers of beads in the assay (biological repetitions) were *n* = 37 in the NHDF-monolayer case and *n* = 36 in the NHDF-intermixed case. The error bars correspond to the standard error of the mean (SEM). (**C**) Distribution bifurcation angles for the monolayer and intermixed fibroblasts seeding conditions. The tabulated values are the mean ± SEM. NHDF-monolayer, *n* = 53; NHDF-intermixed, *n* = 131. The overlap between the histograms is rendered in gray. ‘PDF’ refers to the probability density function.

### Impact of fibroblast distribution on bifurcation angles

In previous literature, the complexity of vascular networks has been typically quantified in terms of the average global characteristics such as the area fraction of the vessels ^48^, lacunarity ^17^ or the so-called branching index, i.e., the linear density of branching points along the vessels ^48^. Here, we additionally focus on *local* morphological features of a branching network, that is on the distribution of the bifurcation angles *ϕ*.

Figure 2C shows the bifurcation angle (*ϕ*) histograms for the advanced stage of vascular networks growth (day 12 of culture) for both ‘intermixed’ and ‘monolayer’ cases. We observe quasi-Gaussian distributions centered around the mean values ϕ^*^ = 55.5+-1.7 deg and ϕ^*^ = 58.3 +-3.2 deg, respectively, with the error estimated as the standard error of the mean. Accordingly, we may conclude that the type of spatial fibroblast distribution has little effect on the observed branching angle. However, in the ‘monolayer’ case, the total number of bifurcation events pooled from all experiments is not excessive, *n* = 53, and it is difficult to draw strong conclusions regarding the measured distribution or the mean value. As we show below, the situation is improved as the size and complexity of the networks increases, which we achieve via increasing the concentration of VEGF in the culture media. At high VEGF concentrations, the number of bifurcation events exceeds *n* = 190 which significantly raises statistical relevance of the results.

### The concentration of VEGF-A-165 determines the onset of angiogenic sprouting, growth dynamics and final morphology of the microvessels

Vascular endothelial growth factors (VEGFs) are essential for the induction of angiogenesis and drive both EC proliferation and migration ^49, 50^. The VEGF family consists of five members: VEGF-A, VEGF-B, VEGF-C, VEGF-D, and placental growth factor (PLGF). The VEGF-A has emerged as the single most important regulator of the blood vessel formation in health and disease; it is essential for embryonic vasculogenesis and angiogenesis, and is a key mediator of neovascularization in cancer and other diseases ^51^. Five isoforms of VEGF-A are known, where the VEGF-A-165 isoform is the one most abundant in humans ^52, 53^.

To characterize the impact of VEGF on EC sprouting dynamics we added increasing concentrations (*C*_VEGF_) of human recombinant VEGF-A-165 into the fibrin bead sprouting assay and quantified the global morphometric parameters of the formed vascular networks. We performed a dose-response titration for a series of VEGF-A-165 concentrations *C*_VEGF_ = [0, 1, 2.5, 5, 10, 25, 50] ng/ml in the medium added to each well in the presence of NHDFs interstitially distributed within the fibrin matrix (the ‘intermixed’ configuration). As in the previous experiments (without the added VEGF), here, we also visualized isolated EC-coated beads at 24 h time intervals over the period of 2 weeks. We observed that, as expected from previous literature ^34, 54-56^, VEGF concentration had a significant impact on the EC sprouting dynamics and overall complexity of the ensuing microvascular networks (Figure 3A and Supp. Figure 3). The EC-coated beads treated with higher *C*_VEGF_ formed larger and more branched vascular networks and started to sprout earlier (Figure 3B). Increasing *C*_VEGF_ from 0 ng/ml to 25 ng/ml led to an approximately 6-fold increase in the final network area *A* and a 3 or 4-fold increase in the final total length *L* (Figure 3Ba and 3Bb). Accordingly, we also observed a 1.5 to 2-fold increase in the final thickness of the branches, *λ* (Figure 3Bc). Increasing *C*_VEGF_ further up to 50 ng/ml did not result in further changes, suggesting saturation of the system at around 25 ng/ml. The observed changes of the area of the mask, *A*_c_ (Figure 3Bd), indicate that the VEGF concentration affected not only the formation of the sprouts but overall had a significant impact on HUVECs proliferation during the assay. In particular, higher concentrations of VEGF resulted in an increased final area of the mask *A*_c_ indicating a more frequent cell proliferation at the monolayer, whereas the lack of the exogenous VEGF (the case 0 ng/ml) led to the decrease in *A*_c_ in the initial stage of the culture. The latter observation could be explained in terms of the arrest of HUVECs growth and/or their apoptosis caused by an insufficient supply of VEGF, resulting in a regression of the vascular network ^57-59^.

**Figure 3.**
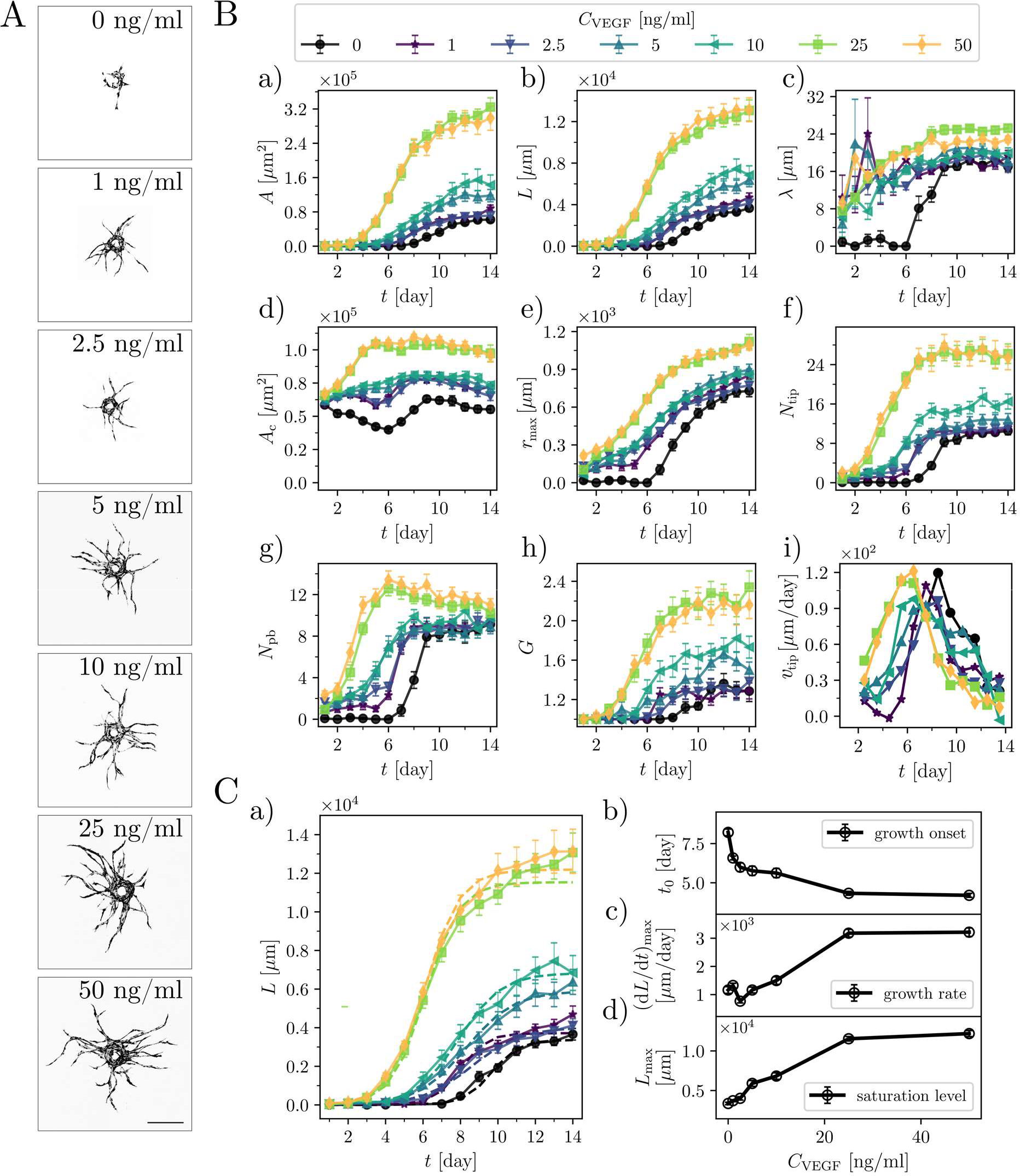
VEGF-A concentration determines the onset of angiogenic sprouting, the rate of growth and the final morphology of the capillary networks. (**A**) Representative images of GFP-tagged HUVEC-coated beads cultured at indicated VEGF-A concentrations, day 10. Scale bar 250 μm. (**B**) Morphometric analysis of sprouting capillary networks at indicated VEGF concentrations in terms of a) total area *A*, (b) total length *L*, (c) average sprout width *λ*, (d) area of the cells coating the bead *A*_*c*_, (e) distance to the furthest tip *r*_*max*,_ (f) number of tips *N*_*tip*_, (g) number of primary branches *N*_*pb*,_ (h) the average number of generations *G* per branch (*G* = log2(*N*_tip_/*Npb*)), and (i) average tip velocity *v*_tip_ (the plot shows moving average with the averaging time of 2 days). The numbers of beads (biological repetitions) taken for the statistics were as follows: *C*_VEGF_ = 0 ng/ml, *n* = 13; *C*_VEGF_ = 1 ng/ml, *n* = 13; *C*_VEGF_ = 2.5 ng/ml, *n* = 14; *C*_VEGF_ = 5 ng/ml, *n* = 14; *C*_VEGF_ = 10 ng/ml, *n* = 12; *C*_VEGF_ = 25 ng/ml, *n* = 14; *C*_VEGF_ = 50 ng/ml, *n* = 12. The symbols in the graphs indicate the mean values and the error bars are the standard error of the mean (SEM). (**C**) The parameters describing the growth process. (a) A comparison between the experimental data *L*(*t*) and fitted curves (a logistic function), see eq. (1), for various *C*_VEGF_. (b) The VEGF-dependence of the onset of growth *t*_0_, (c) the rate of growth (d*L*/d*t*)_max_, and (d) the saturation level *L*_max_. Error bars are the SEM.

The maximum distance *r*_max_ spanned by the networks, the numbers of tips *N*_tip_ and primary branches *N*_pb_ were also significantly elevated for cultures treated with higher *C*_VEGF_ (Figure 3Be-3Bg). Vascular networks exposed to higher VEGF concentrations sprouted earlier and, in general, developed a more complex topology, as reflected by the higher final number of branch generations, *G* = 1+log_2_(*N*_tip_/*N*_pb_) (Figure 3Bh). The increase in *G* was particularly pronounced for cultures treated with *C*_VEGF_ = 25 and 50 ng/ml. This increase in complexity could be attributed, in general, either to the earlier onset of sprouting or to the possible faster linear growth of the sprouts. To verify the latter possibility, we determined the ensemble-averaged linear speed of the tips. We used the formula *v*_tip_ = Δ*L*/(Δ*t N*_tip_), where Δ*t* is the interval between the measurements, that is one day, and Δ*L* is the corresponding increase in the net length *L* of the network (i.e., we assumed that the network grows only at the tips). We found that, independently of the VEGF concentration, the velocity *v*_tip_ as a function of time always developed a maximum shortly after the onset of growth (Figure 3Bi). Importantly, we also observed that the corresponding maximal velocities were of similar magnitude, independently of *C*_VEGF_. Accordingly, we may conclude that the presence of more evolved networks at higher *C*_VEGF_ is associated rather with the earlier onset of sprouting and thus with the overall longer period of growth (in each case extending until saturation at around day 11) rather than with the tip velocity *v*_tip_.

In the following, we turn to a more detail analysis of the evolution of the sprouting networks over time depending on *C*_VEGF_. First, we observe that the two main parameters describing the morphogenesis of a vascular network, i.e., *L*(*t*) and *A*(*t*), exhibit a sigmoidal growth pattern with an initial inactive phase (no sprouts) followed by an exponential growth phase and a final plateau phase. The growth curves *L*(*t*) and *A*(*t*) can be approximated each by a logistic curve of the form:

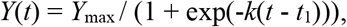

where *Y*_max_ is the saturation level, *k* is the characteristic growth rate in the exponential phase, and *t*_1_ is the inflection point of the sigmoid corresponding to the moment of the fastest growth. From the fitted curves we extract the maximal rate of growth (d*Y*/d*t*)_max_ = *Y*_max_*k*/4 and the onset of growth *t*_0_ = *t*_1_ -2/*k* (root of the tangent at the midpoint). The proposed fits seem to accurately describe the dynamics of EC sprouting (see Figure 3C for the length *L* and Supp. Figure 4 for the area *A*). Some discrepancies (e.g. no obvious plateau phase in some cases) are observed during the final days of the experiment (approximately after day 10 of the assay) and they seem to be more pronounced for higher VEGF concentrations. This is possibly caused by the longer period of extensive growth of the network in these cases, with the growth only starting to saturate at around day 10. We find that the onset of sprout elongation *t*_0_ systematically decreases with *C*_VEGF_ reaching a plateau of *t*_*0*_ = 4 [days] at concentrations *C*_VEGF_ = 25 and 50 ng/ml. This means that increasing *C*_VEGF_ in the range [0, 25] ng/ml expedites sprouting (Figure 3Cb). At low VEGF concentrations (*C*_VEGF_ = 2.5 ng/ml) the maximum rates of growth of the sprout length (d*L*/d*t*)_max_ (Figure 3Cc) and of the sprout area (d*A*/d*t*)_max_ (Supp. Figure 4Ac) are roughly independent of *C*_VEGF_, whereas they are roughly proportional to *C*_VEGF_ in the regime 2.5 ng/mL < *C*_VEGF_ <25 ng/ml and saturate above *C*_VEGF_ = 25 ng/ml. Similar scenarios with a somewhat steadier increase already at the lower *C*_VEGF_ are also observed for the saturation levels *L*_max_ and *A*_max_ (Figure 3Cd and Supp. Figure 4Ad).

**Figure 4.**
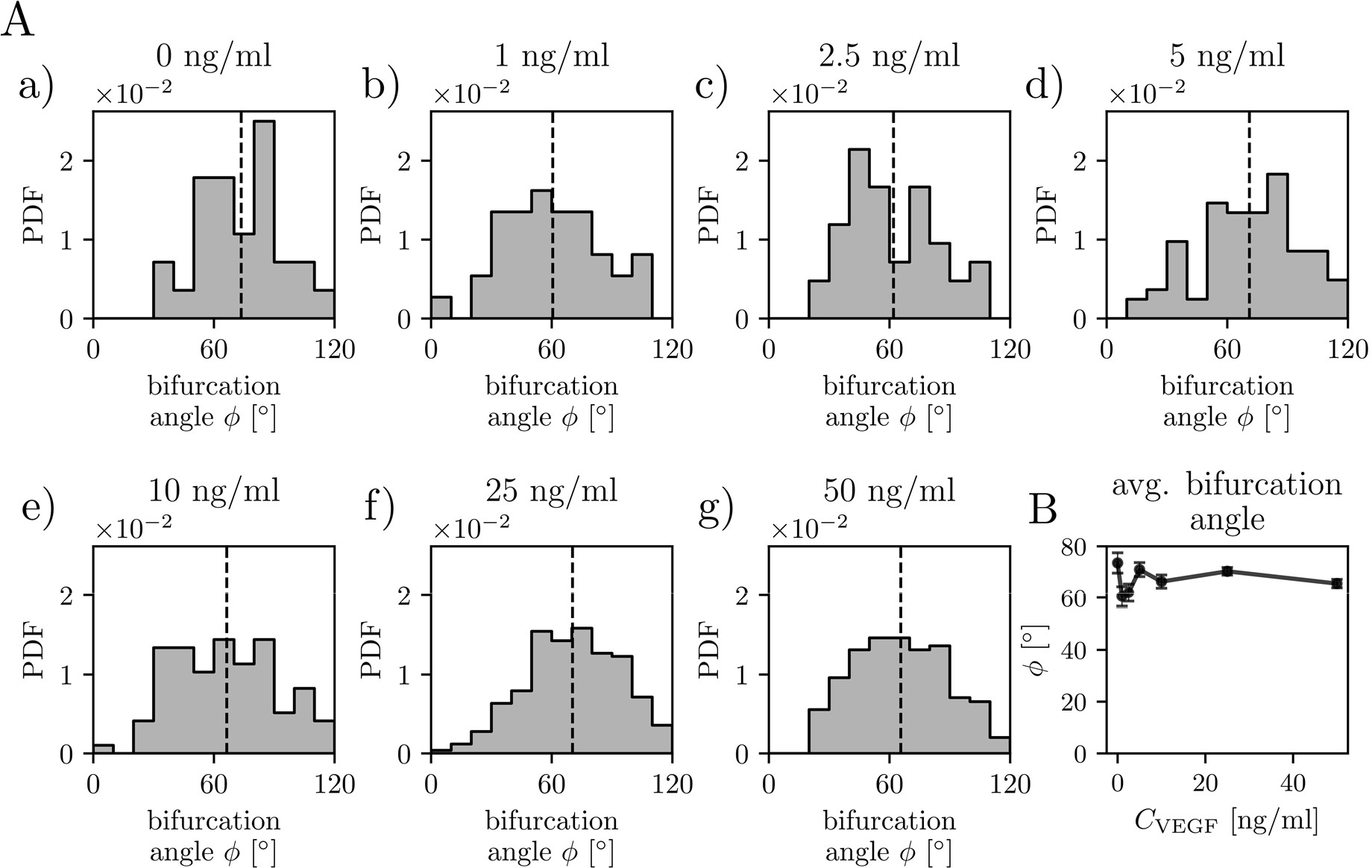
VEGF-A concentration does not affect the distribution of bifurcation angles. (**A**) Analysis of the bifurcation angle distributions for indicated VEGF concentrations at day 12 of culture. The total numbers *n* of the measured angles were: (a) *C*_VEGF_ = 0 ng/ml, *n* = 28; (b) *C*_VEGF_ = 1 ng/ml, *n* = 37; (c) *C*_VEGF_ = 2.5 ng/ml, *n* = 42; (d) *C*_VEGF_ = 5 ng/ml, *n* = 82; (e) *C*_VEGF_ = 10 ng/ml, *n* = 97; (f) *C*_VEGF_ = 25 ng/ml, *n* = 252; (g) *C*_VEGF_ = 50 ng/ml, *n* = 198. The dashed lines show mean values. ‘PDF’ -probability density function. (**B)** The mean bifurcation angle plotted as a function of *C*_VEGF_. The error bars are the standard error of the mean. ‘PDF’ refers to the probability density function.

Finally, we analyze the impact of the VEGF concentration on the distribution of the bifurcation angle *ϕ* (Figure 4) and the segment length *l* (Figure 5). In the case of angle measurements, we first validate our methodology by comparing the results with a manual measurement of the bifurcation angles. Next, we tune the internal parameter of the angle-measurement algorithm (the ‘arm length’, for details see Materials and Methods section, Supp. Figure 5) to best match the manual measurements. With the optimized algorithm, we find that the average bifurcation angle *ϕ*^*^ varies in the range from 61 to 72 deg as *C*_VEGF_ increases from 0 to 50 ng/ml. We observe a shallow minimum at *C*_VEGF_ around 1-2 ng/ml. For higher *C*_VEGF_, *ϕ*\* stabilizes at around 65-68 deg. In the case *C*_VEGF_ = 2.5 ng/ml, we observe two peaks in the bifurcation angle distribution, which might indicate a transition between the effects of small *C*_VEGF_ and the release of bound VEGF from the surrounding fibrin matrix, which becomes important at higher VEGF concentrations ^60^.

**Figure 5.**
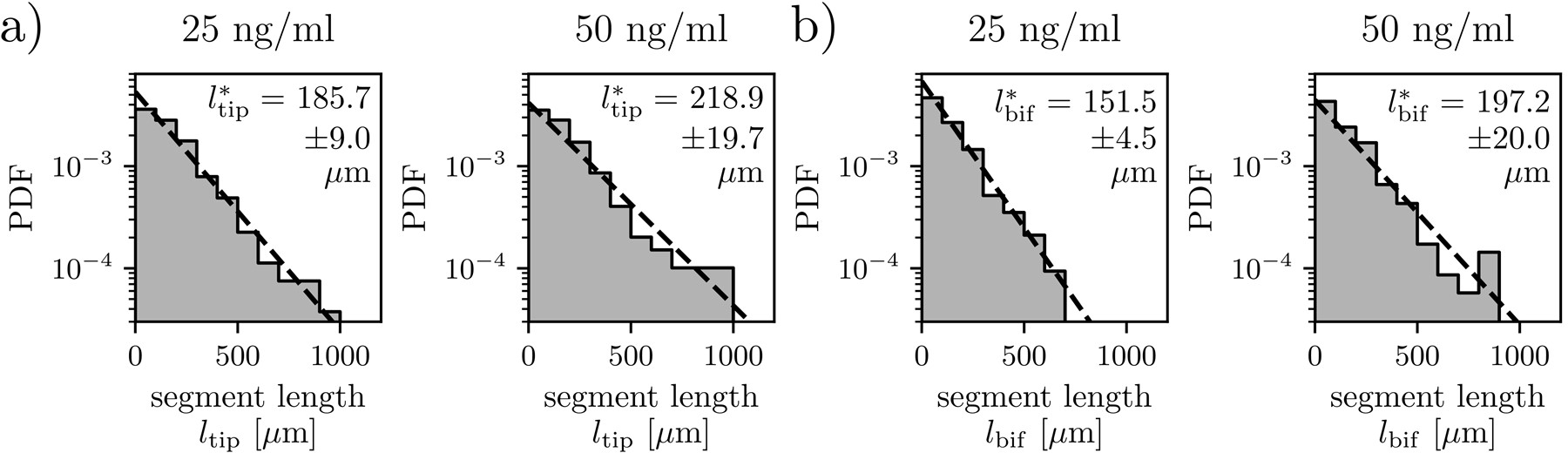
Capillary networks formed at high VEGF concentrations display exponential segment length distributions. Analysis of the distribution of the length of (a) tip segments and (b) bifurcating segments for *C*_VEGF_ = 25 ng/ml and 50 ng/ml. Numbers of identified segments were: (a) *C*_VEGF_ = 25 ng/ml, *n* = 266; *C*_VEGF_ = 50 ng/ml, *n* = 198, (b) *C*_VEGF_ = 25 ng/ml, *n* = 426; *C*_VEGF_ = 50 ng/ml, *n* = 348. The characteristic segment lengths are indicated on the graphs with error being the standard deviation. ‘PDF’ refers to the probability density function.

Considering the distributions *P*(*l*) of the segment length *l*, we also verify the validity of the automated image analysis via comparing with manual image segmentation. In general, we find a reasonable agreement between the automated and manual measurements without any tunable parameters (Supp. Figure 6). The distributions differ only at very small *l*, where apparently the numerical approach overestimates the number of the shortest segments. The overpopulation of the short segments can be attributed to the skeletonization procedure which tends to produce many ‘artificial’ short segments in the regions of high sprout density.

**Figure 6.**
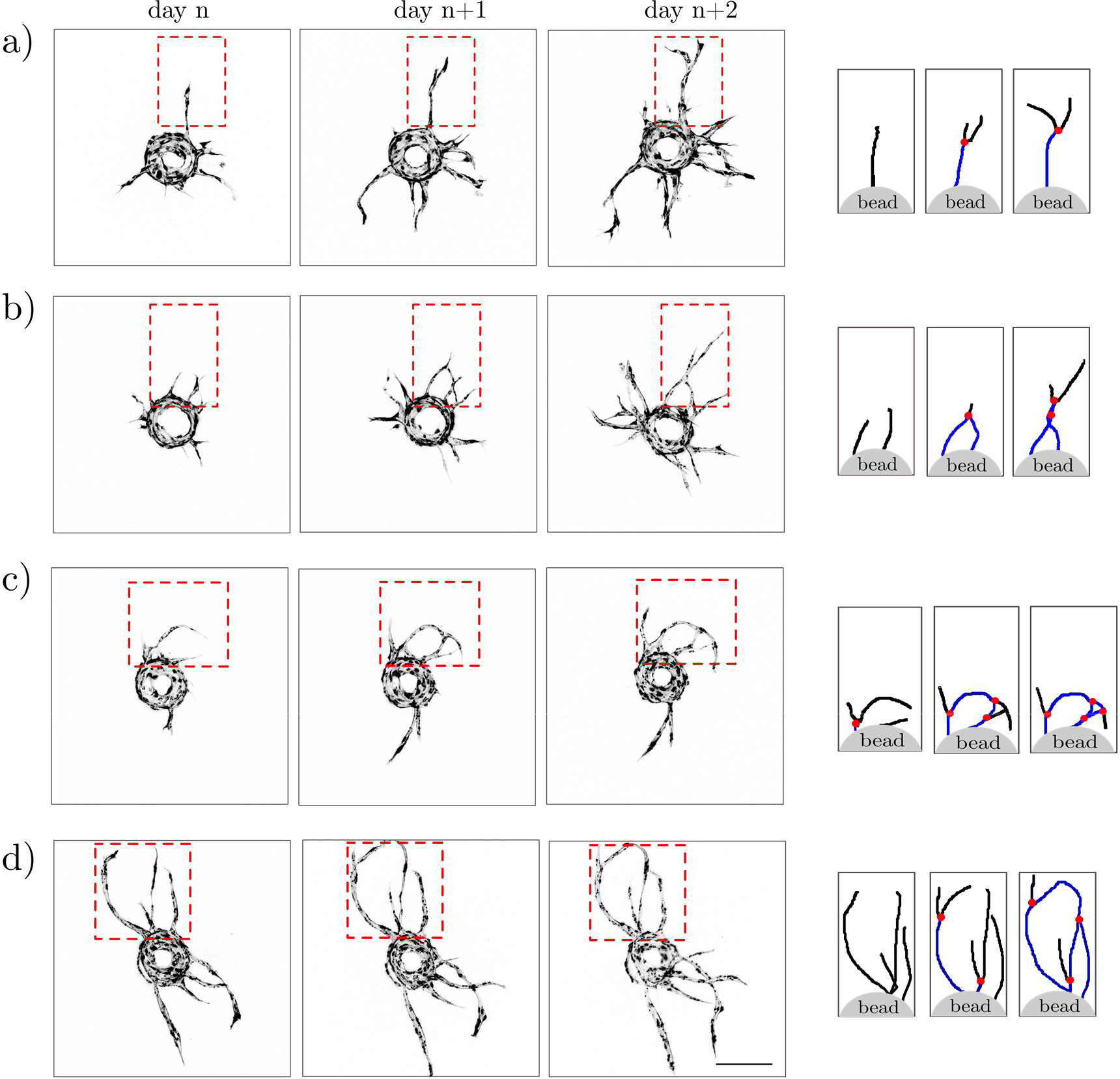
Various scenarios of segment/node formation during development of an EC-bead-sprouting network. Confocal images from three consecutive days of culture and schematic representations of the extracted skeletons of the network with indicated nodes (red dots), tip segments (black lines) and bifurcating segments (blue lines) as would be detected by the algorithm. One can distinguish 3 different scenarios of segment/node formation: (a) bifurcation of a mother branch into two daughter branches, (b) anastomosis (fusion) of two independent branches; here, the anastomosis is followed by another bifurcation, i.e., formation of an X-shaped structure, (c, d) formation of vascular loops caused by sequences of bifurcation and anastomosis events, also including tip-to-tip anastomosis. The scale bar is 250 μm.

Based on results for all studied *C*_VEGF_ we find that at low *C*_VEGF_ the total number of segments is too low to draw statistically meaningful conclusions about the distribution (Supp. Figure 7). Therefore, we limit the more detailed analysis only to highest VEGF concentrations that is the cases *C*_VEGF_ = 25 and 50 ng/ml. In these two cases we observe strong evidence for the exponential decay of *P*(*l*) (Figure 5). The exponentially decaying distributions are observed for both tip segments *P*(*l*_tip_) as well as for the bifurcated segments *P*(*l*_bif_), which is verified via fitting *P*(*l*)∼exp(-*l*/*l*^*^) where *l*^*^ is the average length of a segment. In general, the exponential probability distributions are characteristic of the Poisson process in which the events (bifurcations in this case) occur randomly in space, yet at a constant average spatial density, given by 1/*l*^*^. Another conclusion that can be drawn from Figure 5 is that the average length of a bifurcated segment (*l*^*^_bif_) is significantly shorter than the average length of a tip segment (*l*^*^_tip_), i.e., *l*^*^_bif_ < *l*^*^_tip_. This can be explained by noting that, given sufficient time, an actively growing tip segment will bifurcate and become a bifurcated segment, thus the long ‘tail’ in the distribution builds first in *l*_tip_ distribution and only later becomes visible in *l*_bif_. Finally, no significant difference in *l*^*^_bif_ and *l*^*^_tip_ between the cases with *C*_VEGF_ = 25 ng/ml and *C*_VEGF_ = 50 ng/ml is observed, at least within the statistical error. Noteworthy, we find that the numerical artifact associated with overpopulation of the shortest segments has little effect the fitted values *l*^*^_bif_ and *l*^*^_tip_ (Supp. Figure 6).

## DISCUSSION

During angiogenic sprouting *in vivo*, the growth of the microvessels is driven by local physico-chemical cues, including matrix stiffness and growth factor gradients ^61, 62^ as well as the molecular interplay between VEGF and the Notch signaling pathways ^15, 63, 64^. The coupling of physical and biochemical interactions results in the complexity of the ensuing networks that includes a hierarchical branched structure. In particular, the role of fibroblast distribution in the matrix and the VEGF concentration in the assay have been a subject of intense studies in previous years. Currently, there is a consensus ^32, 33, 47^ that fibroblasts distributed throughout the hydrogel matrix tend to soften the matrix and promote the diffusion of growth factors, which in turn promotes EC sprouting. However, thus far, despite a large body of available literature, the impact of VEGF on the morphology of a vascular network—including the topological network characteristics—has remained an open issue. In particular, the optimal VEGF concentration in the medium has not been clearly identified. In the present study, we report that increasing the VEGF-A-165 concentration, *C*_VEGF_, promotes more rapid development of the networks and leads to more complex vasculatures, an effect which persists up to the saturation level of *C*_VEGF_ ≈ 25 ng/ml VEGF. This is consistent with previous reports ^54, 55, 65-69^ which focused on various biological systems, including *ex-vivo* aortic ring models ^65, 68^, EC spheroids ^69^ and EC-coated microcarrier beads ^54^ as well as vasculature on chip models ^55^, in which VEGF concentrations ranging from app. 30 to 50 ng/ml were observed to induce the fastest growth of vascular networks and promote bifurcations. However, the results of several other studies ^34, 70^ provide contradicting experimental evidence, namely that higher concentrations of VEGF, ranging from 10 ng/ml to 35 ng/ml, result in the fusion of vessels and a reduction of vasculature complexity. For example, Nakatsu and colleagues ^34^ reported a sharp peak of the number of endothelial sprouts (most likely primary branches, but this has not been specified by the authors) in cultures where HUVEC-coated beads were treated with 2.5 ng/ml VEGF, while in assays where VEGF concentration was < 1 ng/ml or > 10 ng/ml the authors observed the formation of approximately one sprout per bead after 7 days of culture. Exposing EC-coated beads to concentrations of VEGF higher than 2.5 ng/ml did not lead to the formation of additional sprouts but resulted in a gradual increase in the sprout width. Part of the confusion may lie in the use of different sources of the HUVEC cells, e.g., different donors and isolation times ^69^ and culture conditions, in particular serum concentration in the culture media, which could partially account for the observed morphometric differences ^71^. In our study, we used commercial HUVEC cells collected from 50 donors and cultured them in a low serum medium, while Nakatsu et al. used freshly isolated HUVEC cells from a single donor and cultured the cells in a high serum medium. The increased serum concentration was previously argued to inhibit the endothelial tube formation ^71^.

The origin of the branched morphology of vascular networks *in vivo* can be related to the proliferative activity of equipotent sprout tips that stochastically bifurcate and randomly explore their environment competing for space ^38^. In the present study, we find that – for relatively high VEGF concentrations – the distribution of lengths of the vascular segments between the bifurcation points, formed by ECs sprouting from EC-coated beads *in vitro*, decays exponentially. This in turn suggests a purely stochastic branching process with a constant bifurcation probability per unit sprout length. Stochastically branching structures are ubiquitous in nature and can be observed both at the level of multicellular organs, such as lungs ^72^, kidneys ^73^, the pancreas ^74^, mammary glands ^75^ or vascular systems ^76^, as well as at the level of single cells such as neurons ^77, 78^ or tracheal cells ^79^. The phenomenon is universal across species ^80^ ranging from prokaryotic organisms ^81, 82^ through invertebrates ^79^ to vertebrates ^72, 77^.

Nevertheless, one should be aware that biological networks are often formed via morphological processes other than branching, yet also resulting in the apparently ‘branched’ structure ^38, 81^ . In our case, for example, we observe that the nodes of the network are formed not only via bifurcation of the sprout tips (Figure 6a), but may also form via (i) sprout fusion (anastomosis), (ii) partial fusion followed by splitting, resulting in the formation of an X-like structure (Figure 6b), or (iii) formation of a vascular loop via tip-to-tip anastomosis (Figure 6c and 6d). The relevance of the abovementioned different morphogenetic scenarios remains an open issue and we leave it for future investigations. Noteworthy, the classification of the nodes of the network with respect to the corresponding scenario of formation would require development of new sprout-tracking algorithms. The fast development of machine-learning (ML) based tools ^83, 84^ promises possible applications of ML also in this field.

Regarding the dependence of the complexity of the networks on VEGF, we observe that increasing the concentration of VEGF leads to a larger number of primary branches and an increase in the total number of bifurcation events. In particular, in the cultures treated with the highest VEGF concentrations (*C*_VEGF_ = 25 and 50 ng/ml), the period of formation of primary branches is followed by the phase of intense branching, resulting in network densification, a process which we do not observe to occur in cultures deprived of exogenous VEGF, or in those treated with low VEGF concentrations (< 2.5 ng/ml). Based on our results we propose that the emergence of more complex morphologies is most likely related to the earlier onset of growth of the network and to the simultaneous emergence of multiple primary sprouts rather than with the faster linear rate of growth of the sprouts. In fact, we found that the maximal speed of the sprout tips is only weakly dependent on VEGF concentration (Figure 3Bi).

Our study suggests that the vascular sprouts grow and bifurcate stochastically, with a constant bifurcation probability per unit length. Based on the observations, the growth process could be likely modeled by a branching random walk ^38^. However, some of the features of the growing vascular networks are not well reproduced by such models. In particular, in the branching random walks, the growing sprout constantly changes its orientation and, as a result, relatively quickly ‘forgets’ its original growth direction. The growth of our vascular networks, although also random to some extent, is nevertheless on the average directed outwards, away from the initial bead, which can be likely related to the direction of local VEGF- and/or nutrient concentration gradient. The directional growth translates into the linear (or even superlinear) dependence of the network size on time (see Figs. 3Bb and 3Bf) shortly after the onset of growth (days 4-8), which is different from the clusters grown in random walk models, the size of which scales sublinearly with time. The random walk models can be adjusted to exhibit radial spreading ^77^ by the introduction of an effective radial force, acting on the growing tips. However, it seems more logical to assume that the growth of tips is guided by the VEGF concentration gradients, which produce a deterministic outward motion on top of stochastic, noise-driven effects. In contrast to the radial force model of Ucar et al. ^77^, such gradients themselves evolve as the sprouts advance into the matrix, competing for the nutrient flux. This makes the vascular growth problem similar to the so-called Laplacian growth models ^85^ in which the structure grows in the direction of the gradient of the diffusive field. In fact, in our experiments, we observe that the average bifurcation angle is typically within the range 65-70 degrees, which is close to the theoretical value of the universal branching angle in the Laplacian growth problems, which is equal to 2π/5, i.e., 72 degrees, at least if the growth takes place in a geometry confined to a plane ^39-42^. Noteworthy, similar values of the branching angles were also reported, e.g., in the quasi-planar retina vasculature ^86^. This suggests that diffusive growth often determines the formation of transport networks. Accordingly, one could use analogies between the growing vascular systems and the evolving non-biological networks, such as river networks or crack patterns, to better understand the dynamics of vascular networks as well as other types of living cellular networks, a consideration which we leave for future studies.

At the end, we shortly discuss the new functionalities of our custom image analysis toolset developed for the purpose of this work. Biological image analysis has enticed the development of numerous tools to aid the interpretation and quantification of experiments, including open-source platforms, such as ImageJ ^87^, extended by FIJI ^88^, CellProfiler ^89^, Vaa3D ^90^, Icy ^91^, and others, as well as commercial tools (such as Imaris, Amira, or Volocity). Particularly popular platforms for angiogenetic analysis are Matlab-based AngioQuant for in vitro assays ^92^ or AngioIQ ^93^ which uses a graphic interface. Carpentier et al. developed Angiogenesis Analyser ^94^ and recently applied it to fibrin-based assays ^95^. These tools offer efficient ways to segment and skeletonize images of angiogenic networks and provide insights into the length and area statistics by resolving junctions and branches. However, they do not include the analysis of bifurcations and branching angle statistics. Our dedicated numerical tool is suited to the format and character of the measured data and tailored to detection of primary branches and outer branches, necessary for the analysis of bifurcations and branching angles. In addition, our software uses Python, and thus benefits from the interoperability and popularity that this language has achieved in recent years. Finally, we note that, with novel tools exploiting the advantages of deep learning coming into play ^96^, we expect that AI will also play an increasingly important role in angiogenesis image analysis.

## CONCLUSIONS

In summary, we have performed high-throughput fibrin-gel angiogenesis bead-assays for the purpose of tracking the evolution of sprouting vascular networks in terms of various statistical morphological/topological characteristics of the networks. To this end, we have developed Phyton-based software for morphometric analysis of vascular networks. We used this tool to demonstrate how the dynamics of growth of HUVEC sprouts depends on the distribution of fibroblasts in the surrounding hydrogel matrix and on the VEGF concentration in the medium. We found that the net length and area of the network, *L*(*t*) and *A*(*t*), or the number of tips *N*_tip_(*t*) develop a characteristic sigmoidal growth pattern consisting of (i) an initial inactive phase, (ii) a rapid growth phase, and (iii) a plateau phase. We also showed that at higher VEGF concentrations, the HUVEC networks start to sprout earlier, produce more initial sprouts (primary branches), and eventually develop longer and more bifurcated networks. Finally, we also studied the distributions of the bifurcation angles and segment lengths in various culture conditions, i.e., depending on the type of the fibroblast distribution (‘monolayer’ or ‘intermixed’) and on the VEGF concentration.

Overall, we have demonstrated the possibility of extracting detailed statistical-topological features of bead-sprouting microvascular networks. Our findings can be of practical relevance in tissue engineering. For example, the HUVEC-coated endothelial ‘seeds’ could be used as dopants to various types of biomaterials or bioinks. The general rules governing the evolution of individual sprouting ‘seeds’ could be then used to determine the optimal seed concentration to warrant efficient vascularization of a biomaterial or a bioprinted construct. Understanding how the external cues affect angiogenic sprouting may also help to optimize tissue repair strategies, e.g., the design of pre-vascularized wound dressings. Moreover, in the future, the use of custom-generated hydrogel microcarriers—instead of the commercially available polystyrene microbeads—could open the possibility of co-encapsulation of various cell types inside the beads together with ECs, e.g. various tumor lines, supporting cells, etc., and allow development of miniaturized, vascularized cancer models for applications, e.g., in drug testing.

## MATERIALS AND METHODS

### Cell culture

GFP-HUVECs (Angio-Proteomie, Boston, MA; catalog no. cAP-0001GFP) were cultured in EBM-2 medium supplemented with EGM-2 bulletkit (Lonza, Basel, Switzerland; catalog no. CC-3156 & CC-4176) and were used at passages 3 through 5. NHDF (Promocell, Heidelberg, Germany; catalog no. C-12302) were cultured in Dulbecco’s modified Eagle medium (DMEM) (Thermo Fisher Scientific, Waltham, MA, USA; catalog no. 10566016) supplemented with 10% fetal bovine serum (FBS), 4.5 g/l glucose, Glutamax, 1% penicillin-streptomycin. NHDF were used between passages 2 and 7. All cells were cultured in 5% CO_2_ at 37 °C humidified atmosphere and media were replaced every 2 days.

### Fibrin bead-sprouting assay

Coating of the beads with EC was performed as described previously ^97^ with small modifications. Briefly, HUVECs-GFP were mixed with 265μm diameter monodispered polystyrene superparamagnetic microcarrier beads (microParticles GmbH, Berlin, Germany; catalog no. PS-MAG-AR111) at the concentration of approximately ∼ 500 cells per bead in a small volume of warm EGM-2 medium and placed in the incubator for 4 hours at 37°C and 5% CO_2_, gently shaking the tube every 20 min. After 4 hours beads were transferred to culture flask with fresh EGM-2 medium and placed over night in the incubator at 37° C and 5% CO_2_. The following day beads were gently washed with EGM-2 medium, resuspended in freshly prepared 2.5 mg/ml fibrinogen solution (Sigma-Aldrich, St. Louis, MO, USA; catalog no. 341573), mixed with 0.625 units of thrombin (Sigma-Aldrich, St. Louis, MO, USA; catalog no. T4648) and seeded at one bead/well in 24-well plate placing the bead centrally in the well. Fibrin/bead solution was allowed to clot for 5 minutes at room temperature and then at 37° C and 5% CO_2_ for 30 minutes. 1mL of EGM-2 medium was added to each well. NHDF at a concentration of 25 000 cell/well were either layered at the top of the clot or added to the fibrin/bead solution before seeding the beads. Medium was changed every day. Human recombinant VEGF-165 (Stemcell technologies; Saint Egrève, France; catalog no.78073) was used at the indicated concentrations.

### Immunofluorescence staining

Fibrin blocks with NHDF cells were fixed with 4% paraformaldehyde (PFA), and blocked with blocking buffer (2% bovine serum albumin [BSA], 2% normal goat serum, and 0.5% Triton X-100 in PBS). Actin-488 conjugated antibody (Sigma-Aldrich, St. Louis, MO, USA; catalog no. ABT1485-AF488) was applied in blocking buffer.

### Image acquisition and processing

EC-coated beads were imaged every 24 hours for 14 consecutive days using Nikon A1 confocal microscope (Nikon Instruments, Inc, Melville, NY, USA) equipped with PLAN APO 10×/0.45 objective. Images were collected using NIS-Elements Advanced Research software (Nikon Instruments, Inc, Melville, NY, USA) in the nd2 format, which are 16-bit single-channel images with respective metadata. One experiment was recorded as a set of successive frames, where a single frame had several slices in the z-direction. The resolution of images is 1.25 μm/pixel. The images were first max-pooled on z-directional slices (taking the maximum intensity value across the stack) and treated with a Gaussian blur with a kernel of size 11 pixels. To distinguish cells from the background, segmentation was performed with the threshold based on the average intensity multiplied by 1.17. A filling algorithm was used to eliminate holes with perimeter smaller than 200 pixels (250 μm), and the largest connected component was taken. The centrally located polystyrene bead (mask) was then detected by a top-hat transform algorithm. To focus the analysis on the geometrical characteristics of the growing sprouts the central bead was removed using the previously detected mask. The processed images were subsequently skeletonized using an algorithm from the scikit-image package. Connectivity of the resulting skeletonized network was then determined by identifying branching points and branches (segments). Finally, outer branches (with Horton index 1) shorter than 50 pixels (62.5 μm) were pruned, except for those connected to the mask.

### Geometrical and topological characteristics of the sprouting network

Based on the segmented pictures, skeleton and mask, we computed the geometrical characteristics of the sprouting network (*A*_c,_ *A, L, r*_max,_ *N*_pb,_ *N*_tip,_ *λ, G*). Moreover, for each experiment, we detected the bifurcations points, that is the nodes in the graph with three outgoing segments and measured three angles associated with each bifurcation point. The angles were measured between the straight lines connecting the bifurcation point and a selected point on each of the outgoing branches. Because of the finite thickness of the sprouts, the process of skeletonization resulted in branches being slightly curved towards the branching point. Accordingly, if one selected the points for the angle measurement very close to the bifurcation point, then the computed bifurcation angle would appear larger than in the case with the points selected at a larger distance from the bifurcation. We decided to select the points at 60 pixels (approx. 75 μm) from the bifurcation point (calculated along the sprout) on each branch as in this case we obtained the results that best matched the manual measurements (see Supp. Figure 5).

It is important to note that out of the three angles adjacent to a bifurcation point, one needs to select the actual bifurcation angle, that is the angle between the two *daughter* branches. In general, based on manual segmentation of several images, we found that the bifurcation angles are typically between 30 and 120 deg. Therefore, in the algorithm, we took the smallest of the three angles as the supposed bifurcation angle. We note that this method has a limitation in the sense that it cannot yield bifurcation angles larger than 120 deg. Also, the smallest angle at a selected node may as well be associated with anastomosis of the vessels rather than bifurcation. In order to exclude the angles corresponding to anastomosis events we checked whether the bisector of the selected smallest angle pointed inwards or outwards with respect to the polystyrene bead (the center of the mask). Only the cases with the bisector pointing outwards (i.e., with the maximum angle of 90 deg between the bisector and the vector connecting the center of the bead and the node) were classified as bifurcations.

Finally, the fitting of the exponentially decaying function to the PDF of branch lengths *P*(*l*) was done using semi-logarithmic scale. That is, we fitted a linear function with a negative slope to the dataset ln(*P*(*l*)) versus *l*. To this end we used the least squares method. The fitting of a sigmoid function from eq. (1) to the experimental data *L*(*t*) (Figure 3Ca) and *A*(*t*) (Supp Figure 4a) was done using non-linear least squares method.

## Supporting information

Electronic Supplementary Information

## ACKNOWLEGMENTS

Preparation of this article was supported by Sonatina grant (no. 2020/36/C/NZ1/00238 awarded to K.O.R.), Opus grant (no. 2022/45/B/ST8/03675 awarded to J.G.) and Sonata grant (no. 2018/31/D/ST3/02408 awarded to M. L.) from the Polish National Science Center (NCN). The authors thank Leon Jurkiewicz for assistance in data analysis.

