## Supplementary material for "Topological evolution of sprouting vascular networks: from day-by-day analysis to general growth rules": Electronic Supplementary Information

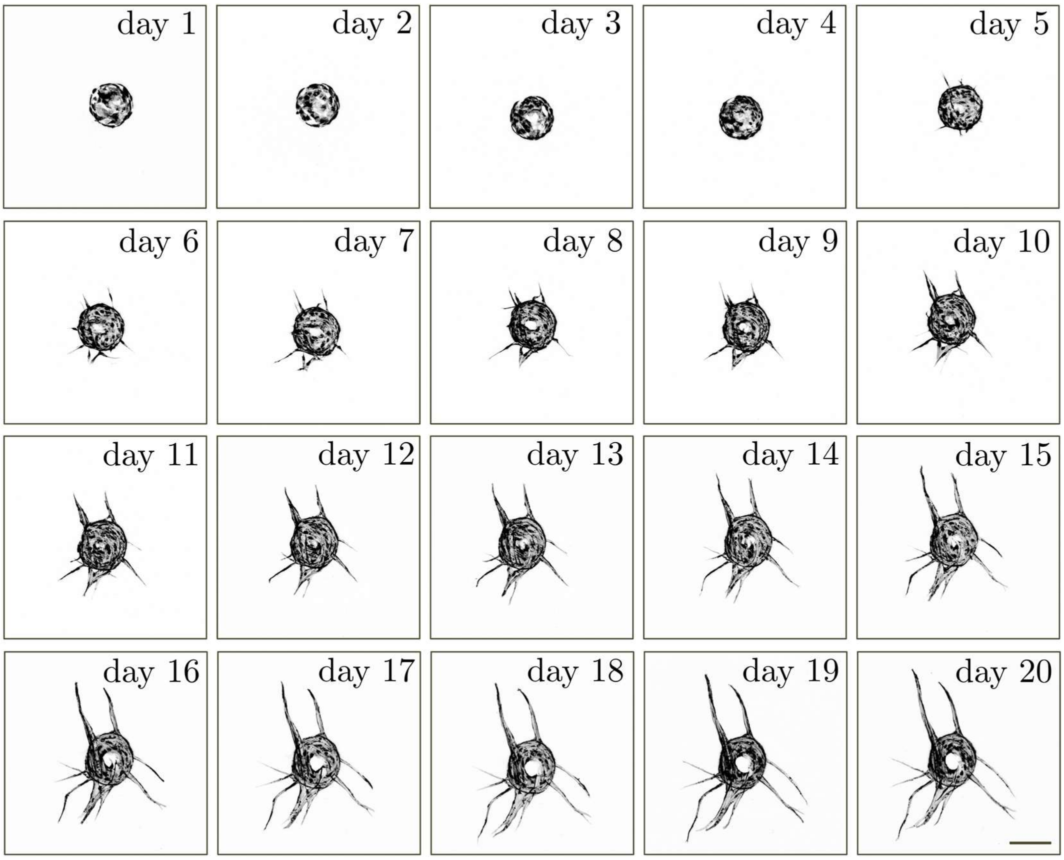


**Supplementary Figure 1. Long term culture of GFP-tagged HUVEC-coated beads.** Representative images of EC-coated bead acquired at 24h intervals for 21 consecutive days. Cells were visualized with GFP. Scale bar 250 µm.

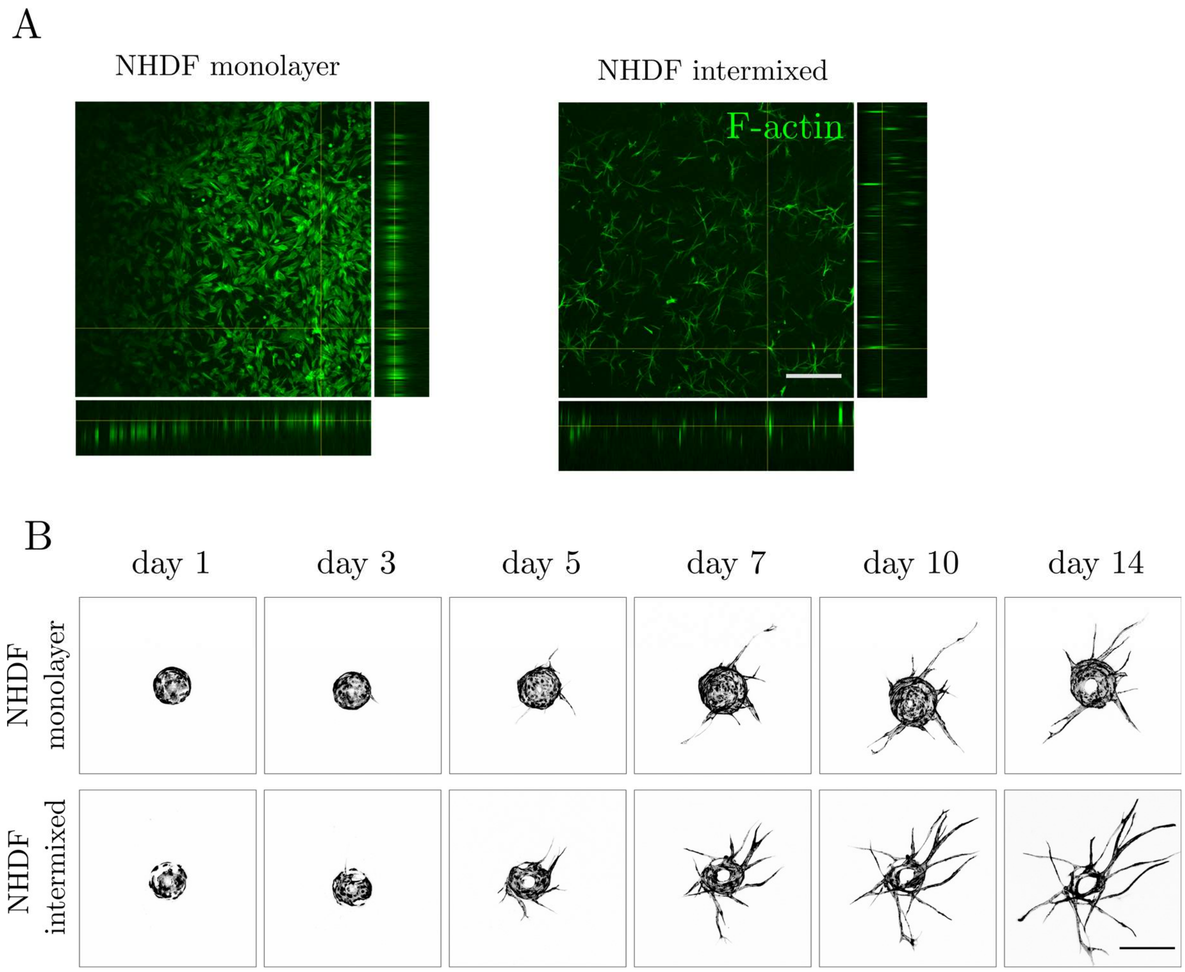


**Supplementary Figure 2. Morphology of capillary networks formed by GFP-tagged HUVEC-coated beads cultured at different NHDF seeding conditions (A)** Confocal images and orthogonal projections showing NHDF distribution when cultured as a monolayer at the top of the fibrin clot (left panel) or intermixed within the fibrin gel (right panel). Cells were visualized with Alexa-488 conjugated F-actin antibody. Scale bar 250 µm. (**B**) Representative images of single GFP-tagged HUVEC-coated beads cultured in monolayer and intermixed condition at day 1, 3, 5, 7, 10 and 14. Scale bar 250 µm.

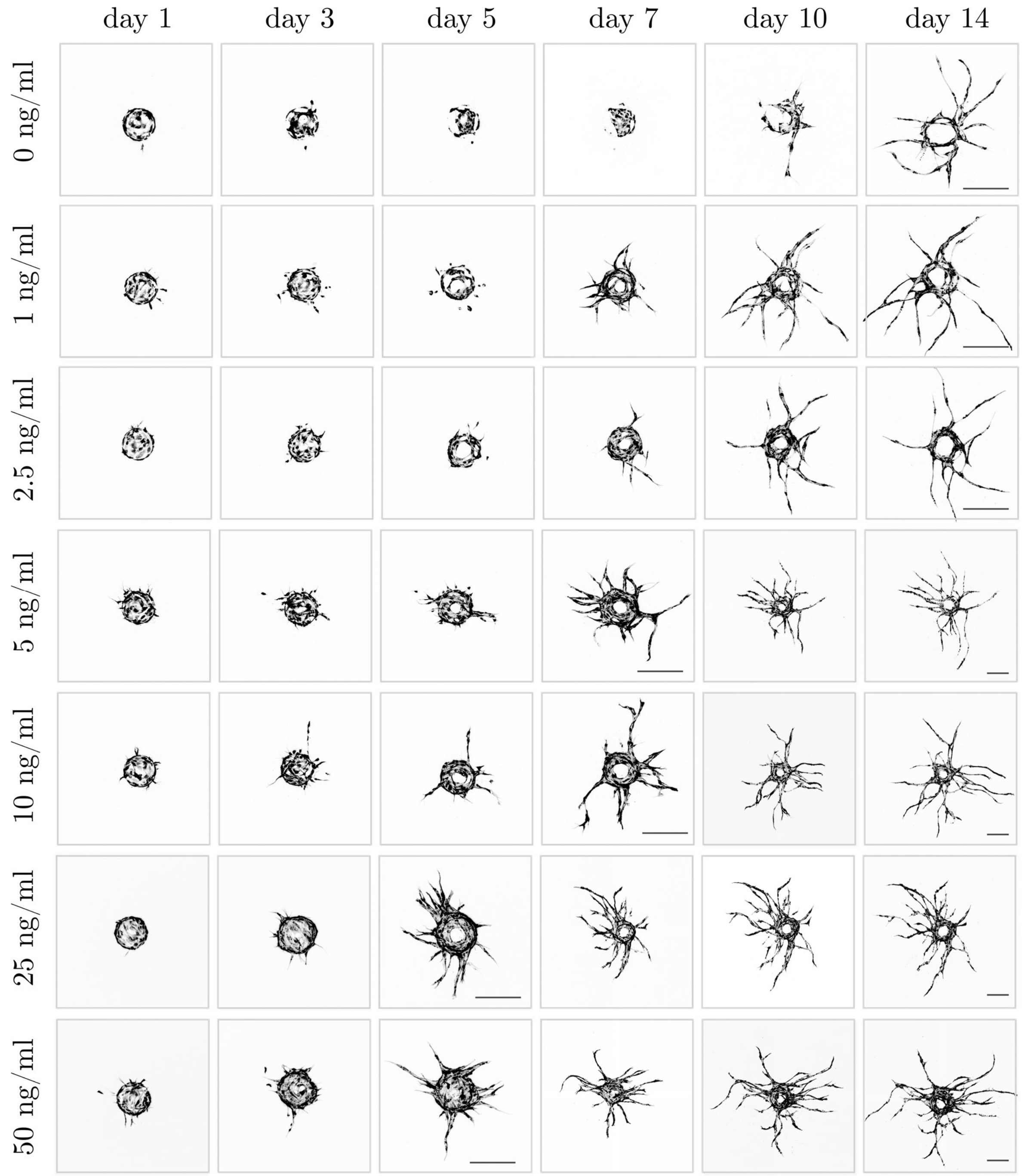


**Supplementary Figure 3. Morphology of capillary networks formed by GFP-tagged HUVEC-coated beads cultured at different VEGF concentration.** Representative images of single GFP-tagged HUVEC-coated beads cultured at indicated VEGF concentrations at day 1, 3, 5, 7, 10 and 14. Scale bar 250 µm.

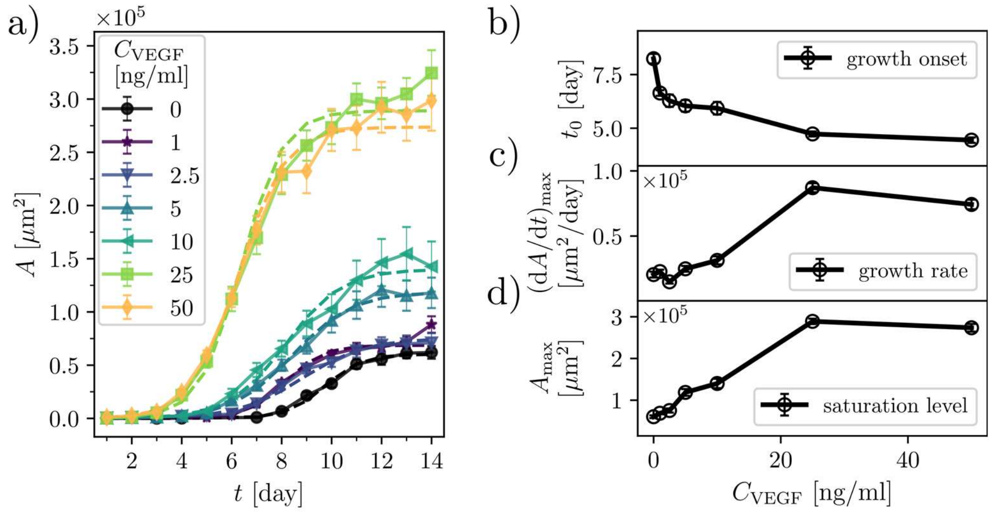


**Supplementary Figure 4. Characterization of the dynamics of EC sprouting at different VEGF concentrations.** (a) A comparison between the experimental data $A(t)$ and the fitted curves (a logistic function), see eq. (1), for various $C_{\text{VEGF}}$. (b) The VEGF-dependence of the onset of growth $t_0$, (c) the rate of growth $(\mathrm{d}A/\mathrm{d}t)_{\max}$, and (d) the saturation level $A_{\max}$. Error bars are the SEM.

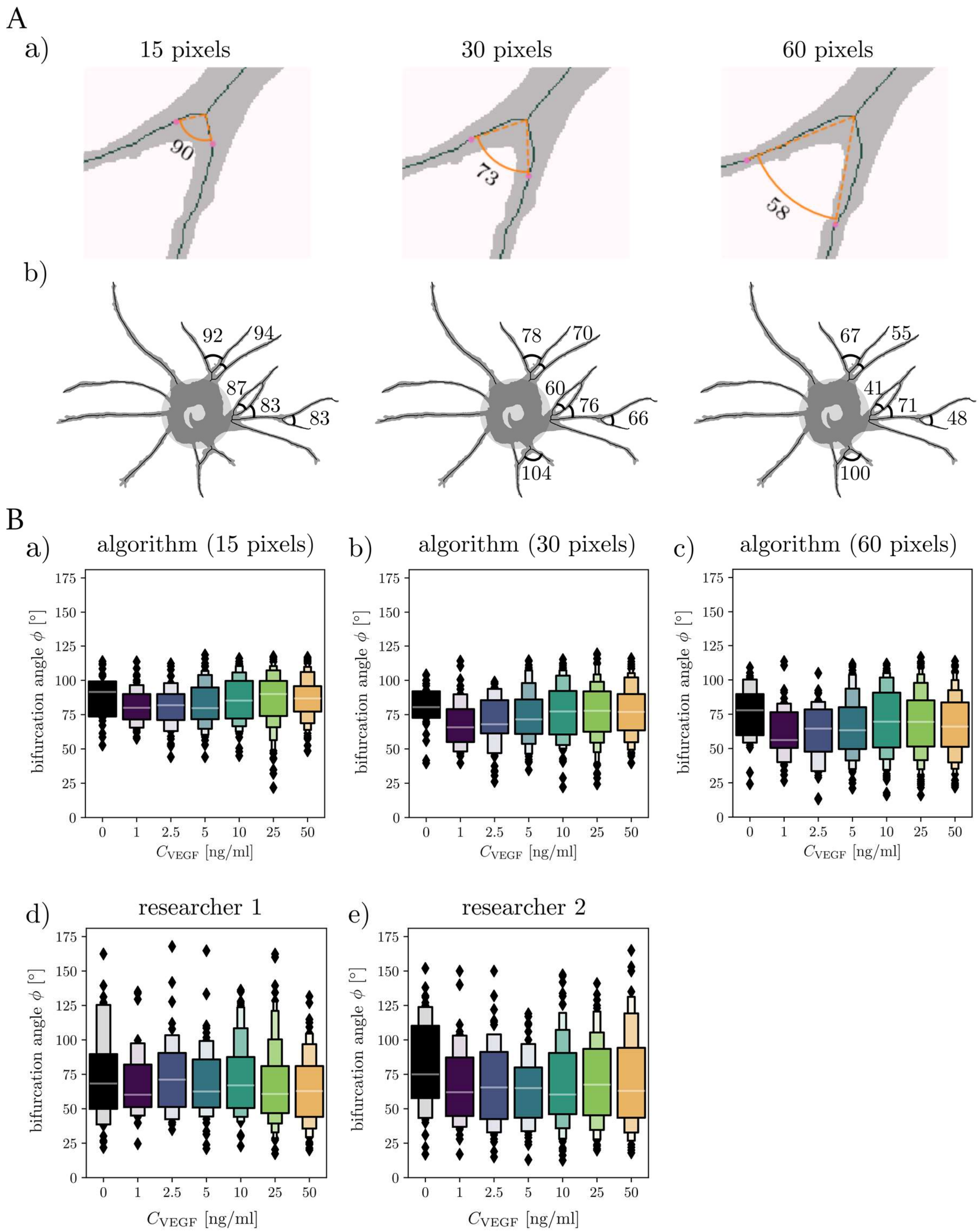


**Supplementary Figure 5. Values of the branching angles depend on the localization of the points between which the angle is measured.** (**A**) Representative images of the (a) single bifurcating branch and (b) EC-sprouting beads showing the values of bifurcation angles (in degrees) with the angle arms defined by the points along the daughter branches located at a distance of 15, 30 or 60 pixels from the bifurcation point. Angle located on the bottom of the bead on the left image in (b) exceeded 120 degrees and was excluded by the software. (**B**) Comparison of (a-c) automated (algorithmic) and (d-e) manual measurements (researcher 1 and 2) of the bifurcation angles at indicated VEGF concentrations at day 14 of culture. Manual measurements were performed in a blind manner by two independent researchers. The numbers of the bifurcation angles: (a) detected by the algorithm in the case 15 pixels: $n$ = [29, 50, 51, 86, 112, 243, 201]; (b) the case 30 pixels: $n$ = [30, 50, 52, 86, 112, 250, 207]; (c) the case 60 pixels: $n$ = [32, 48, 52, 83, 114, 246, 204], (d) researcher 1: $n$ = [44, 40, 48, 47, 108, 78, 64]; researcher 2: $n$ = [35, 34, 39, 50, 74, 138, 93].

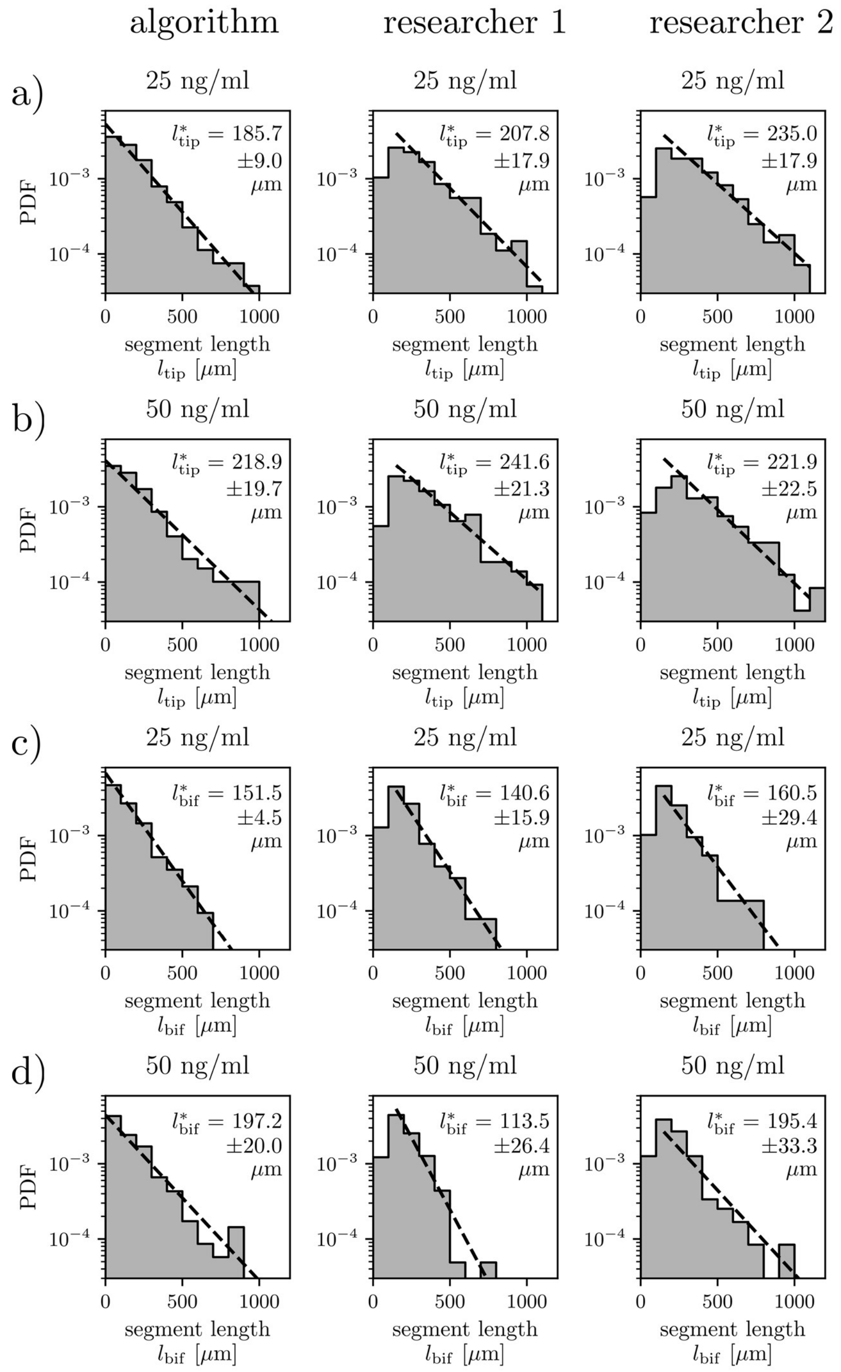


**Supplementary Figure 6. Distribution of segment lengths at high VEGF concentrations. (A)** Comparison of automated (algorithmic) and manual measurements (researcher 1 and 2) of (a, b) the tip segment lengths and (c, b) the bifurcating segment lengths for (a, c) $C_{\mathrm{VEGF}}$ = 25 ng/ml and (b, d) $C_{\mathrm{VEGF}}$ = 50 ng/ml. Characteristic segment lengths are indicated on the graphs with the error being the standard deviation. Manual measurements were performed in a blind manner by two independent researchers. The numbers $n$ of detected segments in the three cases—algorithm, researcher 1, researcher 2—were, respectively: (a) $n$ = 266, $n$ = 271, $n$ = 281, (b) $n$ = 198, $n$ = 217, $n$ = 240, (c) $n$ = 426, $n$ = 257, $n$ = 147, (d) $n$ = 348, $n$ = 205, $n$ = 119. ‘PDF’ refers to the probability density function.

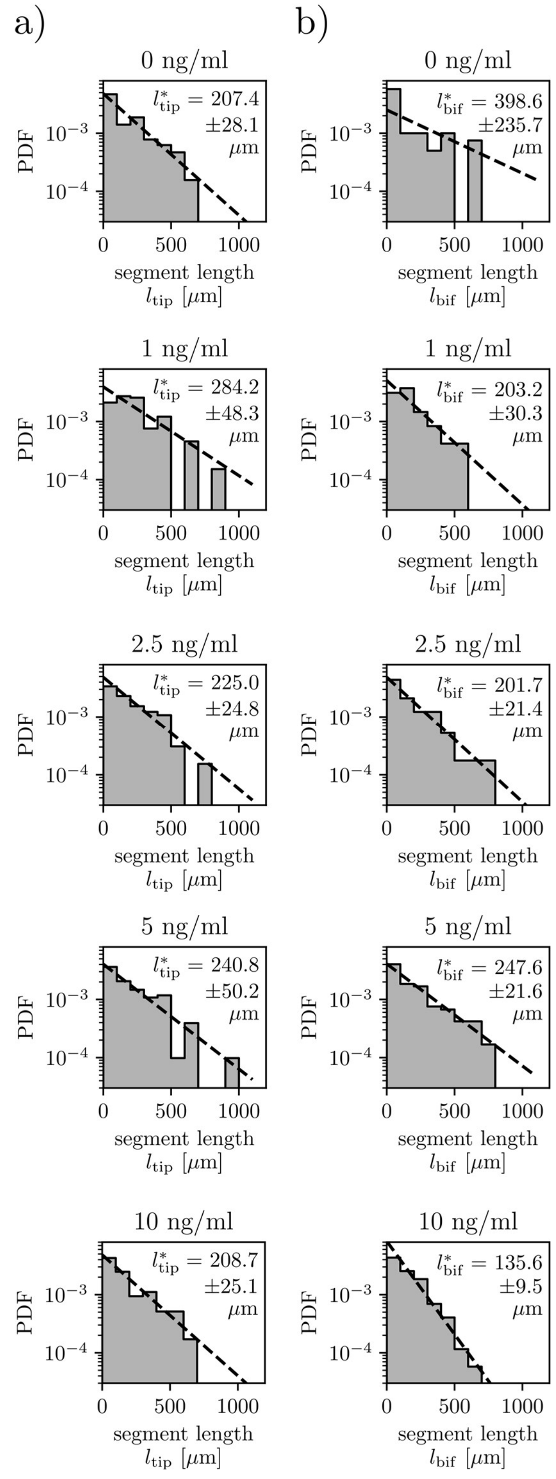


**Supplementary Figure 7. Distribution of segment lengths at lower VEGF concentrations.** Analysis of distributions of (a) the tip segment lengths and (b) the bifurcating segment lengths at indicated VEGF concentrations. Numbers $n$ of segments detected by the algorithm were: (a) $C_{\mathrm{VEGF}} = 0$ ng/ml, $n = 64$; $C_{\mathrm{VEGF}} = 1$ ng/ml, $n = 66$; $C_{\mathrm{VEGF}} = 2.5$ ng/ml, $n = 65$; $C_{\mathrm{VEGF}} = 5$ ng/ml, $n = 102$; $C_{\mathrm{VEGF}} = 10$ ng/ml, $n = 117$, (b) $C_{\mathrm{VEGF}} = 0$ ng/ml, $n = 40$; $C_{\mathrm{VEGF}} = 1$ ng/ml, $n = 48$; $C_{\mathrm{VEGF}} = 2.5$ ng/ml, $n = 57$; $C_{\mathrm{VEGF}} = 5$ ng/ml, $n = 119$; $C_{\mathrm{VEGF}} = 10$ ng/ml, $n = 173$. Characteristic segment lengths are indicated on the graphs with the error being the standard deviation. 'PDF' refers to the probability density function.
